# Peptide-to-protein data aggregation using Fisher’s method improves target identification in chemical proteomics

**DOI:** 10.64898/2026.02.02.702201

**Authors:** Hezheng Lyu, Hassan Gharibi, Zhaowei Meng, Bohdana Sokolova, Xuepei Zhang, Roman A. Zubarev

**Affiliations:** Division of Physiological Chemistry I, Department of Medical Biochemistry and Biophysics, Karolinska Institutet, 171 65 Stockholm, Sweden; Biomotif AB, 183 48 Täby, Sweden; Chemical Proteomics, Swedish National Infrastructure for Biological Mass Spectrometry (BioMS), Stockholm, Sweden; Chemical Proteomics Unit, Science for Life Laboratory (SciLifeLab), Stockholm, Sweden

## Abstract

Protein-level statistical tests in proteomics aimed at obtaining *p*-value are conventionally made on protein abundances aggregated from peptide data. This integral approach overlooks peptide-level heterogeneity and ignores important information coded in individual peptide data, while protein *p*-value can also be obtained by Fisher’s method of combining peptide *p*-values using chi-square statistics. Here we test this latter approach across diverse chemical proteomics datasets based on assessments of protein expression, solubility and protease accessibility. Using the top four peptides ranked by their *p*-values consistently outperformed protein-level analysis and avoided biases introduced by inclusion of deviant peptides or imputation of missing peptide values. Fisher’s method provides a simple and robust strategy, improving identification of regulated/shifted proteins in diverse proteomics assays.

## Introduction

Differential expression analysis between two groups of samples is the most common type of analysis performed in proteomics. One of the most important steps is the determination of *p*-value, which is the *a priori* probability for protein abundance to differ by the observed or higher value. The Student’s t-test applied to data from several replicate analyses is commonly used for such purpose. In proteomics, the *p*-values obtained for individual proteins need to be corrected for multiple hypotheses using, e.g., Bonferroni or Benjamini-Hochberg correction^1^. Often, few (if any) statistically significant proteins remain after stringent correction, even though many protein abundances seem to change. This gives rise to the false negative problem in, e.g., drug discovery, where multiple drug targets need to be identified and characterized.

Part of the problem is in the suboptimal data processing. Search engines commonly used in bottom-up proteomics, such as Proteome Discoverer^2^, MaxQuant^3^, and Mascot^4^, extract first peptide abundances and then aggregate them into protein abundance. This is done by the algorithms that are often not transparently documented. Sometimes, the abundances of peptides belonging to the same protein are simply summed together. The peptides attributed to a given protein by mistake or those with highly fluctuating abundances (e.g., due to post-translational modifications, PTMs) will also contribute to the result, reducing its statistical significance. If the peptide data are missing in some replicates, they are frequently imputed by arbitrary values. The aggregated protein abundance can therefore be a mixture of reliable and questionable peptide data, which leads to higher (poorer) *p*-values than justified.

There are approaches to filter peptide data before aggregating them into proteins, such as the Diffacto technique^5^. These approaches are based on investigating peptide behavior in different samples through covariation analysis and rejecting the deviating peptides. Such techniques work particularly well when several non-replicate samples (e.g., obtained by different treatments) are present in the dataset, and require substantial bioinformatic efforts. A simpler and more straightforward approach could be to use Fisher’s formula for merging *p*-values of independent series of measurements^6^. As unique peptides in proteomics are statistically independent entities measured in several replicates, a *t*-test can be performed for each peptide. Thus we hypothesized that it should be possible to integrate with advantage these peptide data into a protein *p*-value via Fisher’s formula.

To test this hypothesis, we reanalyzed using different peptide-to-protein strategies several chemical proteomics datasets obtained in our lab by a variety of proteomics techniques. This includes the datasets assessing the changes in protein expression (FITExP^7^ and ProTargetMiner^8^), and solubility (PISA^9^ and OPTI-PISA^10^). In order to determine which variation of Fisher’s approach, if any, is optimal, we designed the Figure of Merit (FoM) based on the combined ranking of 244 known targets of 115 drugs. As the results clearly demonstrated the superior performance of Fisher’s-based technique over the conventional approach, the optimal analytical strategy was validated on the datasets assessing the protease accessibility of protein domains (AFDIP^11^ and HOLSER^12^).

## Methods

### Data processing and abundance normalization

Each dataset was composed of several raw LC-MS/MS data files of tryptic peptide mixtures with tandem mass tag (TMT) multiplexing. The data were processed with either MaxQuant (MQ) or Proteome Discoverer (PD) analysis programs, or both. In the latter case, both MQ and PD-obtained results were used in further analysis. Peptide-level outputs were used for Fisher’s analysis, while protein-level outputs served for comparison with Fisher’s results. Peptides and proteins assigned to contaminant proteins or reversed sequences were removed from the results, as were peptides not uniquely assigned to one protein. Peptides were grouped by protein IDs, and peptide abundances were normalized to the total ion count within each TMT channel. For expression (FITExP) and solubility (PISA) datasets in the method development part of the study, fold changes were calculated directly from protein abundances. For the validation datasets obtained by partial digestion (AFDIP^11^ and HOLSER^12^), fold changes were calculated from the abundances of top four peptides.

In protein-level analysis, molecules identified with modified peptides only were filtered away. In both analysis types, identifications with more than one missing value among the biological replicates (n≥3) per condition (treatment or control) were removed, as well as proteins with fewer than two peptides. In the PISA-Expression dataset, any remaining missing values were imputed using the average normalized abundance in other replicates. Protein lists from both analysis types were cross-validated, and only proteins present in both lists were retained to ensure fair comparison.

### Statistical testing

For each peptide and protein, two-tailed, unpaired Student’s t-test was performed comparing normalized abundances of treated versus control samples. Fold changes were calculated as the ratio of the median (among the replicates) abundances in treated samples to control samples. For PISA, expression, and OPTI-PISA data, only peptides with consistent fold-change direction were retained. Benjamini-Hochberg (BH) multiple testing correction was applied, and the resulting adjusted *p*-values were used in downstream analysis.

In peptide-level analysis, peptides belonging to the same protein ID were ranked by their *p*-value, fold changes or a composite score defined as -log10(*p*-value) × |log2(fold change)|. In Fisher’s method, either all peptides or the top N peptides (N = 2, 3, … 6) for each protein were used. When N exceeded the number of available peptides for a given protein, missing values were handled in one of the two ways: (i) using only the available *p*-values (“no imputation”), or (ii) imputing the average *p*-value of the existing peptides for the same protein.

### Fisher’s method of p-value aggregation

For each protein, *X*^*2*^ value was calculated from the *p*-values of *N* selected peptides:

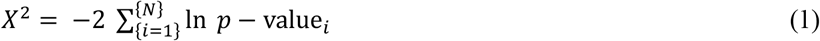

The combined protein *p*-value was computed as the one-tailed probability of observing a *X*^*2*^ value equal to or greater than the observed *X*^*2*^. This was done using the pchisq() function in R.

### Figure-of-Merit for evaluation of strategies

The basic requirement was that analytical strategies that rank known targets higher (lower Rank value) should produce larger FoM values. For each drug but staurosporine, FoM was calculated as:

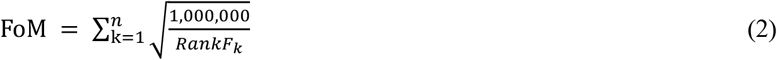

Here, RankFk - ranking factors of identified known targets (from DrugBank^13^), and n is the number of such targets. Ranking factors were calculated as follows: for each protein, a score was computed by combining the fold change of protein abundance with its associated *p*-value, obtained either from BH-corrected t-tests (protein-level analysis) or from Fisher’s method (peptide-level analysis). The score was defined as -log10(*p*-value) × |log2(fold change)|. In the HOLSER^12^ and AFDIP^11^ datasets, fold changes of proteins were calculated either as the average fold changes or center of gravity shifts of the top N peptides, where the direction (positive or negative) was determined by the majority of selected peptides. Proteins were sorted by their scores in the descending order, and the obtained ranks were then rescaled to a uniform range of 1–8000 to enable comparison of datasets with different proteome depths. These rescaled ranks represented the ranking factors in (2). For staurosporine, a broad-spectrum kinase inhibitor with hundreds of kinases known to bind the compound, we counted the number of kinases within the top 100 ranked proteins, which served as the FoM score.

Lastly, the FoM values for individual datasets were renormalized for the same N to a 1–10 scale, and the rescaled values were summed together to obtain the final Score for a given strategy.

### Datasets used for method evaluation

The proteomics datasets are listed in Table 1, and all known targets of drugs used in this study are given in Supplementary Table 1.

**Table 1.**
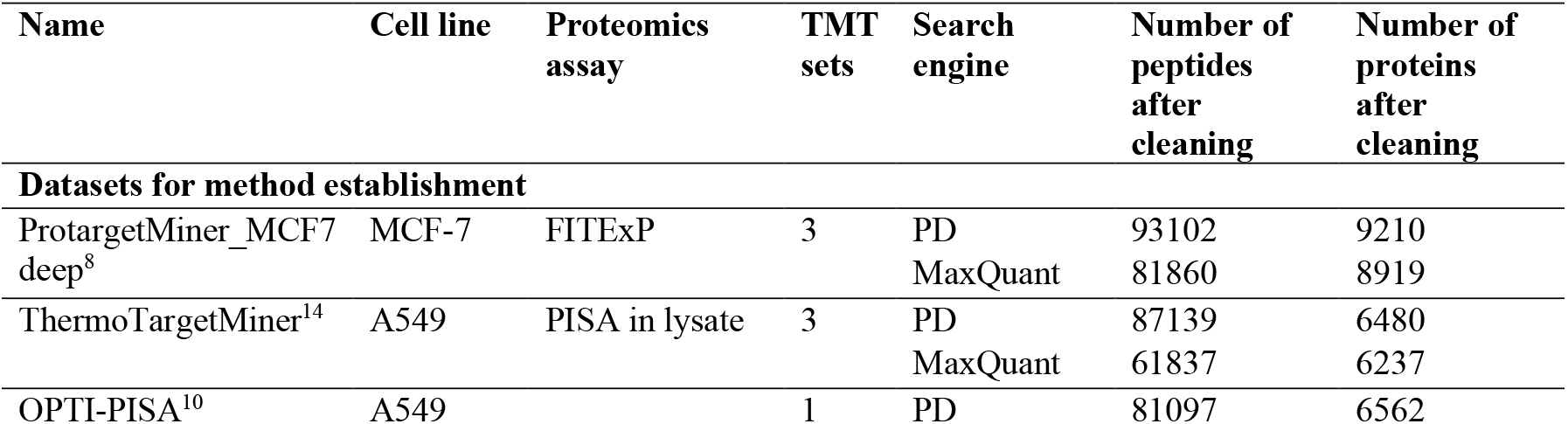
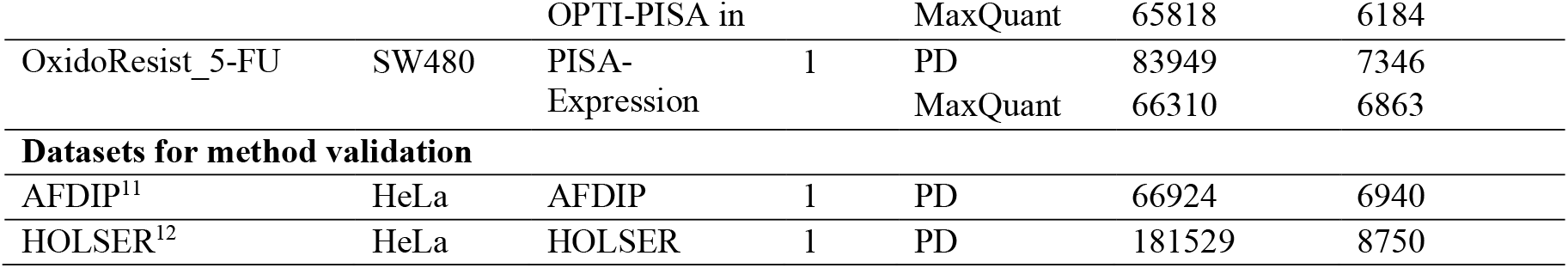
Datasets used for the Fisher’s method in proteomics data analysis.

The first dataset used for testing Fisher’s-based approach consisted of expression (FITExP) proteomic data from drug-treated MCF-7 cells extracted from the ProTargetMiner database^15^. Briefly, MCF-7 cells were exposed for 48 h to IC_50_ concentrations of nine anticancer compounds representing diverse mechanisms of action. For FoM calculations, we used the known targets of bortezomib (PSMB1 and PSMB5) and raltitrexed (TYMS and FPGS). The second dataset was a solubility (PISA^9^) assay from the ThermoTargetMiner database^14^ and originated from A549 cell lysate treated with different drugs against lung cancer. For FoM determination, we used the known targets of vorinostat (targets: HDACs), everolimus (target: MTOR) and olaparib (targets: PARP1, PARP2, and AKR1C3). The third dataset was an automated PISA assay (OPTI-PISA^10^) obtained from A549 cell lysates treated with 10 μM methotrexate (MTX, target: DHFR), staurosporine (targets: kinases), or ganetespib (targets: HSP90s). The last dataset comprised the combined PISA-Expression^16^ proteomics profile of 5-fluorouracil (5-FU)-resistant versus 5FU-sensitive SW480 cells, with the 5FU targets of TYMS used for FoM calculations.

The datasets used for validation of the optimal Fisher’s-based strategy encompass data on protein domain accessibility to trypsin in the AFDIP^11^ and HOLSER^12^ techniques. In both cases, known targets of MTX (target: DHFR), staurosporine (targets: kinases), and rapamycin (targets: MTOR and FKBPs) were used for FoM determination.

## Results and Discussion

As the majority of proteins in most datasets were identified with six or more peptides (an example is shown in Figure 1A), the range of tested *N* values was limited to six peptides. In the discovery part of the study, we found by comparing the FoM-based Scores (Figure 1B and Supplementary Table 2) that for all N, ranking modes and imputation strategies Fisher’s method provides significantly better performance compared to conventional protein-level analysis. That is, known targets universally received on average better ranking when protein *p*-values in chemical proteomics data were obtained by aggregating peptide *p*-values using Fisher’s formula.

**Figure 1.**
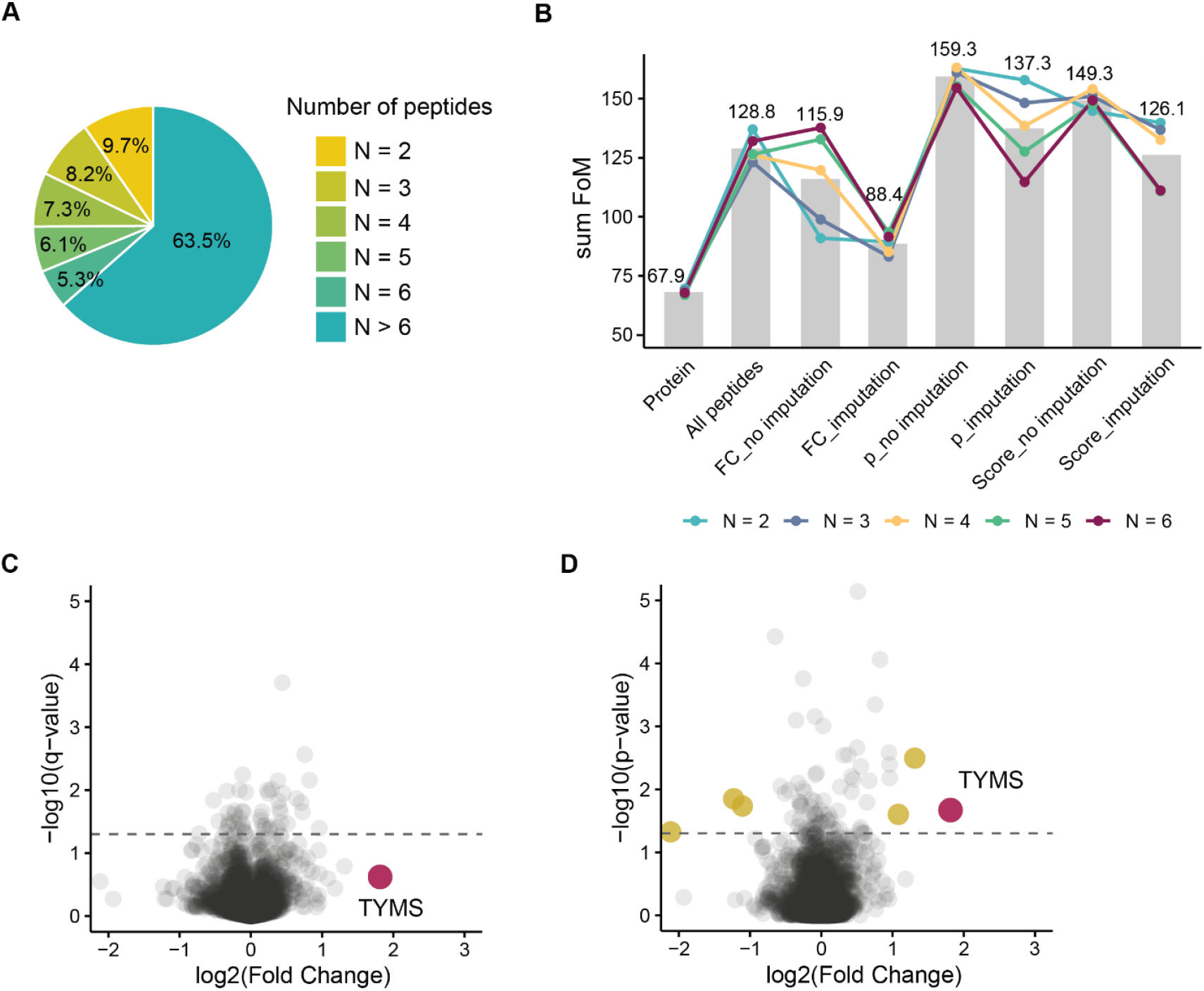
A. Distribution of peptide coverage of proteins in the PISA-Expression dataset of 5-FU resistant versus sensitive SW480 cells. B. Scores for different Fisher’s-based approaches. C. Raltitrexed’s target TYMS in the ProTargetMiner dataset did not pass the significance threshold based on protein expression changes. D. Fisher’s method applied to the top four peptides per protein ranked by Student’s t-test *p*-value identified the known target of raltitrexed, TYMS, highlighted in purple. Other possible candidates are shown in yellow.

A closer look showed that incorporating imputed *p-*values when the number of available peptides was lower than N consistently reduced the analysis quality. Ranking peptides by their *p*-value obtained from *t-*test was found to be the best, followed by ranking by a combination of *p*-value and fold change. Note that this ranking refers to the selection of peptides for Fisher’s analysis, while after protein *p*-value calculation the final protein ranking was done by combined score as described in the Methods.

Including in the analysis all available peptides for each protein was somewhat problematic because proteins with a larger number of peptides tended to yield artificially small aggregated *p*-values, introducing bias toward highly covered proteins. Therefore, detailed investigation was performed on the effect of N on the FoM values. Using top four peptides ranked by Student’s *t*-test *p*-value yielded the most accurate target identification.

As an example, in the ProTargetMiner deep-proteome dataset, the raltitrexed target TYMS ranked 9^th^ in the protein-level analysis and did not even reach the *p*-value threshold of 0.05. In contrast, applying Fisher’s method to the top four peptides ranked by the *p*-value elevated TYMS to the top position by the combined score (Figure 1C and Figure 1D).

To validate the developed Fisher’s-based approach, we applied it to detecting ligand-induced conformational changes by partial digestion (AFDIP^11^ and HOLSER^12^). Unlike the expression and solubility proteome profiling (FITExP^7^ and PISA^9^), where peptides belonging to the same protein behave the same way, in partial digestion only peptides related to drug binding change their abundance. Therefore, we expected that Fisher’s approach with limited N will be particularly suitable in these techniques compared to the traditional protein-based fold-change approach. A case in point is presented in Figure 2A and Figure 2B, where none of the known rapamycin targets (MTOR and FKBPs, shown in purple) ranked among the top five proteins in the target candidate list obtained by conventional analysis. The top protein, CBR1, has no established connection to MTOR signaling. In contrast, using Fisher’s approach to the top four peptides with the lowest *p*-values as well as their fold changes as protein fold change revealed significant allosteric shifts for FKBP2 and FKBP3 (Figure 2C and Figure 2D).

**Figure 2.**
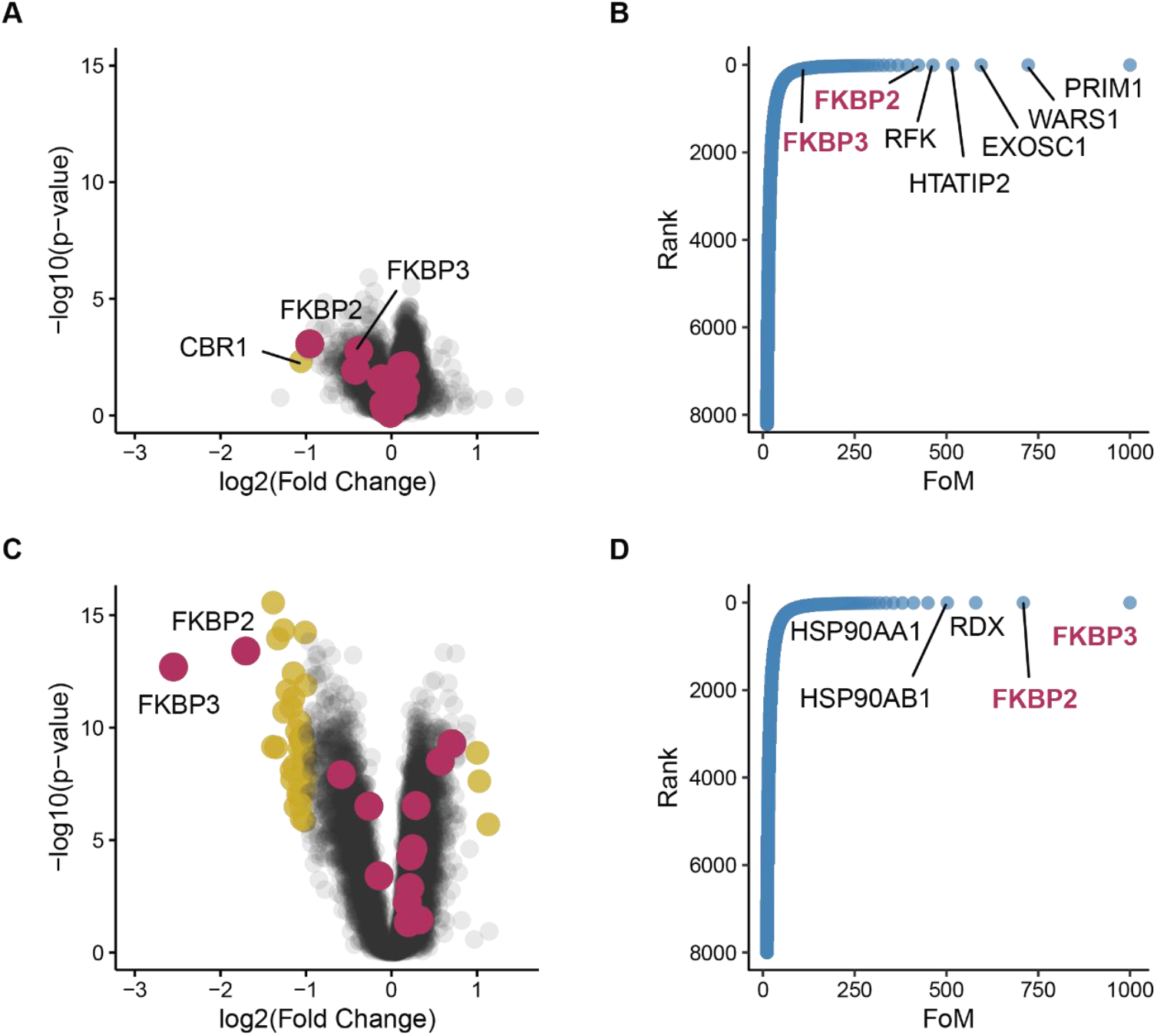
Comparison of rapamycin target identification in the HOLSER dataset using either protein abundance data (A and B) or Fisher’s method applied to the four peptides with the smallest *p*-values without imputation (C and D). In the volcano plots (A and C), significance cut-offs for fold changes are ±1, and for *p*-value is 0.05. Significant proteins are shown in yellow, while known rapamycin targets (FKBPs and MTOR) are highlighted in purple. Waterfall plots (B and D) show proteins being ordered by their FoMs.

A similar advantage was observed in the AFDIP peptide-level dataset^11^. Here, center of gravity differences ΔCoG between1 treated and control 8-h digestion curves were quantified. While conventional approach did not produce any significant candidates after B.-H. correction, applying Fisher’s method to calculate protein *p*-value from top four peptides and averaging their ΔCoG values to represent ΔCoG of a protein correctly identified four known rapamycin targets, FKBP2, FKBP3, FKBP4, and MTOR on the top of the ranked protein list (Figure 3).

**Figure 3.**
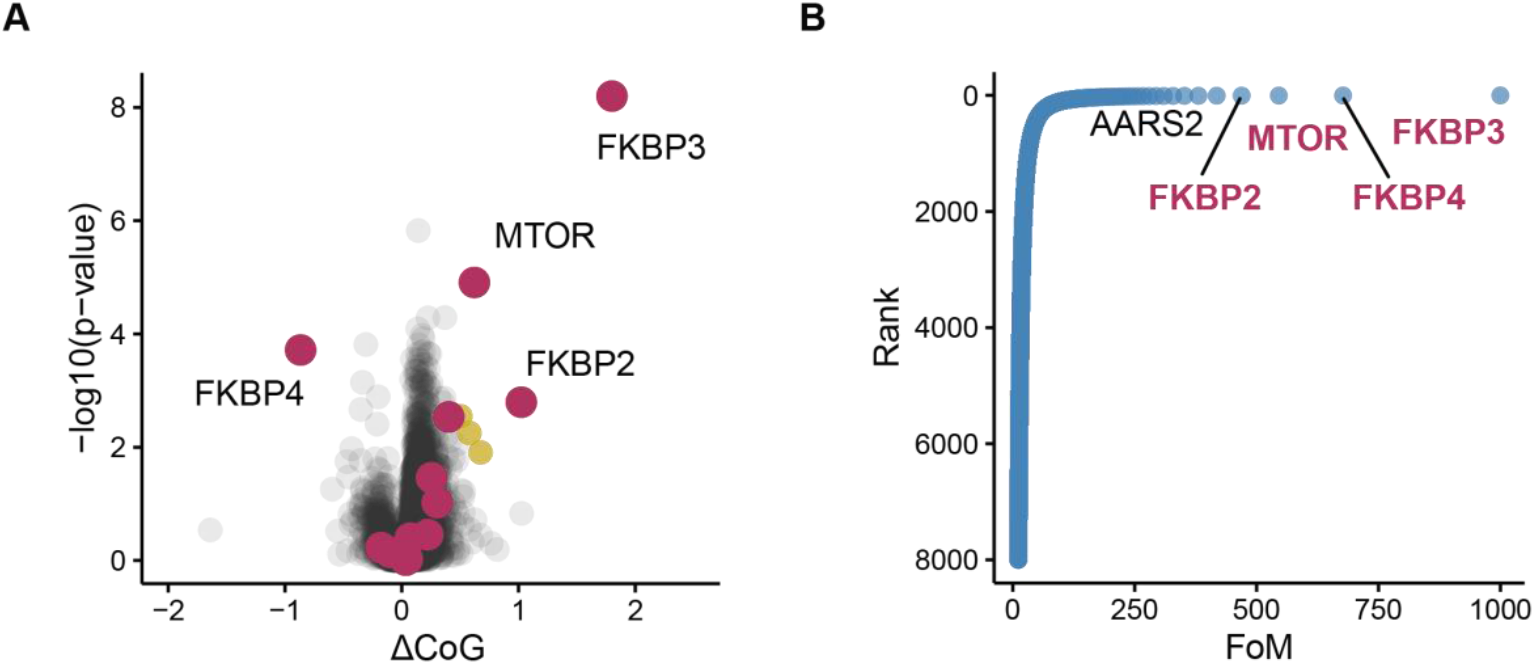
Application of Fisher’s method for rapamycin target identification in AFDIP data. Fisher’s method was applied using the top four peptides per protein, ranked by *p*-value and analyzed without missing value imputation. A. Volcano plot. B. Ranking of protein targets with known rapamycin targets FKBP2, FKBP3, FKBP4 and MTOR occupying the top four positions.

## Discussion

The results presented above demonstrate that aggregating peptide-level statistical evidence provides a more sensitive and robust strategy for protein-level inference in chemical proteomics than conventional protein abundance-based analysis. By operating directly on peptide-level *p*-values, Fisher’s method preserves heterogeneity among peptides belonging to the same protein and avoids dilution of true signals caused by noisy or uninformative peptide measurements. This effect was consistently observed across datasets representing protein expression, solubility, and ligand-induced conformational changes, indicating that the improvement is not assay-specific but reflects a general analytical advantage.

Although Fisher’s method formally assumes independence among combined tests, peptide-level measurements originating from the same protein are not strictly independent in all cases. Nevertheless, the consistent performance gains observed across diverse datasets, search engines, and proteomics platforms suggest that the approach is empirically robust to moderate peptide dependence. Importantly, restricting the aggregation to a limited number of top-ranked peptides further mitigates potential bias arising from peptide redundancy and shared technical variation.

A key observation of this study is that including all peptides associated with a protein can introduce systematic bias, as proteins with higher peptide coverage tend to yield artificially small aggregated *p*-values. Limiting the analysis to the top N peptides ranked by statistical significance effectively controls this bias while retaining sensitivity to true effects. Across all evaluated datasets, using the top four peptides provided the most reliable balance between robustness and sensitivity, highlighting peptide selection as a central component of accurate protein-level statistical inference.

Handling missing peptide-level information was found to be equally critical. Imputation of missing *p*-values consistently reduced analytical performance, likely due to artificial inflation of statistical evidence. In contrast, restricting Fisher’s aggregation to observed peptide-level *p*-values yielded more accurate and reproducible protein ranking, emphasizing the importance of conservative missing-value handling in peptide-based statistical frameworks.

The advantages of peptide-level *p*-value aggregation were particularly pronounced in partial-digestion-based assays, where only a subset of peptides proximal to ligand-binding sites is expected to respond to treatment. Under such conditions, conventional protein-level aggregation is inherently suboptimal, whereas Fisher’s method effectively captures localized peptide responses. As a downstream data analysis strategy, the proposed approach requires no changes to experimental design or quantification workflows and can be readily integrated into existing proteomics pipelines.

## Conclusion

In this study, we demonstrated that Fisher’s method provides a simple yet powerful statistical framework for integrating peptide-level information into protein data in proteomics analysis workflow. By combining *p*-values of top peptides rather than relying on statistical analysis of aggregated abundances of all peptides, this approach effectively captures peptide-level heterogeneity, filters away unreliable peptide data and enhances the detection of biologically relevant targets in chemical proteomics. Across multiple datasets representing protein expression, solubility, and conformational changes, Fisher’s method consistently outperformed conventional protein-level analyses. These findings establish Fisher’s method as a robust and generalizable strategy for improving identification of significantly changing proteins across diverse proteomics platforms.

## Data Availability

The raw mass spectrometry proteomics data and search/quantification results are available with the ProteomeXchange Consortium via the PRIDE repository under accessions: ProtargetMiner PXD013134, ThermoTargetMiner PXD054158, OPTI-PISA PXD050241, PISA-Expression PXD071434, HOLSER PXD069119 and AFDIP PXD061498.

## Code availability

All analysis scripts, R code, and processing pipelines used in this study are available on GitHub at https://github.com/annlyu96/PeptideFisher.

## Acknowledgement

This work was supported by the Swedish Research Council (grant number 2021-05223), Cancerfonden (22 1967 Pj) and EU (consortia ARIADNE VIBE and ALLODD).

## Reference

(1) Silva, T. S.; Richard, N. Visualization and Differential Analysis of Protein Expression Data Using R. Methods Mol Biol 2016, 1362, 105–118. 10.1007/978-1-4939-3106-4_6.

(2) Orsburn, B. C. Proteome Discoverer—A Community Enhanced Data Processing Suite for Protein Informatics. Proteomes 2021, 9 (1), 15. 10.3390/PROTEOMES9010015.

(3) Cox, J.; Mann, M. MaxQuant Enables High Peptide Identification Rates, Individualized p.p.b.-Range Mass Accuracies and Proteome-Wide Protein Quantification. Nat Biotechnol 2008, 26 (12), 1367–1372. 10.1038/NBT.1511.

(4) Mascot search engine | Protein identification software for mass spec data. https://www.matrixscience.com/.

(5) Zhang, B.; Pirmoradian, M.; Zubarev, R.; Kall, L. Covariation of Peptide Abundances Accurately Reflects Protein Concentration Differences. Mol Cell Proteomics 2017, 16 (5), 936. 10.1074/MCP.O117.067728.

(6) Sir Ronald Fisher, B. A.; DSc Ames, frs; Calcutta, L. Statistical Methods for Research Workers THIRTEENTH EDITION-REVISED.

(7) Chernobrovkin, A.; Marin-Vicente, C.; Visa, N.; Zubarev, R. A. Functional Identification of Target by Expression Proteomics (FITExP) Reveals Protein Targets and Highlights Mechanisms of Action of Small Molecule Drugs. Scientific Reports 2015 5:1 2015, 5 (1), 1–9. 10.1038/srep11176.

(8) Saei, A. A.; Beusch, C. M.; Chernobrovkin, A.; Sabatier, P.; Zhang, B.; Tokat, Ü. G.; Stergiou, E.; Gaetani, M.; Végvári, Á.; Zubarev, R. A. ProTargetMiner as a Proteome Signature Library of Anticancer Molecules for Functional Discovery. Nat Commun 2019. 10.1038/s41467-019-13582-8.

(9) Gaetani, M.; Sabatier, P.; Saei, A. A.; Beusch, C. M.; Yang, Z.; Lundström, S. L.; Zubarev, R. A. Proteome Integral Solubility Alteration: A High-Throughput Proteomics Assay for Target Deconvolution. J Proteome Res 2019, 18 (11), 4027–4037. 10.1021/acs.jproteome.9b00500.

(10) Meng, Z.; Saei, A. A.; Lyu, H.; Gaetani, M.; Zubarev, R. A. One-Pot Time-Induced Proteome Integral Solubility Alteration Assay for Automated and Sensitive Drug-Target Identification. Anal Chem 2024, 96 (48), 18917–18921. 10.1021/acs.analchem.4c05127.

(11) Sokolova, B.; Gharibi, H.; Jafari, M.; Lyu, H.; Lovera, S.; Gaetani, M.; Saei, A. A.; Zubarev, R. A. Above-Filter Digestion Proteomics Reveals Drug Targets and Localizes Ligand Binding Site. bioRxiv 2025, 2025.03.11.642584. 10.1101/2025.03.11.642584.

(12) Zhang, X.; Sokolova, B.; Meng, Z.; Gharibi, H.; Gaetani, M.; Zubarev, R. A. High-RatiO PartiaL ProteolysiS with CarriER Proteome (HOLSER) Enables Global Structure Profiling and Site-Resolved Elucidation of Ligand–Protein Interactions. bioRxiv 2025, 2025.07.11.664381. 10.1101/2025.07.11.664381.

(13) Knox, C.; Wilson, M.; Klinger, C. M.; Franklin, M.; Oler, E.; Wilson, A.; Pon, A.; Cox, J.; Chin, N. E. L.; Strawbridge, S. A.; Garcia-Patino, M.; Kruger, R.; Sivakumaran, A.; Sanford, S.; Doshi, R.; Khetarpal, N.; Fatokun, O.; Doucet, D.; Zubkowski, A.; Rayat, D. Y.; Jackson, H.; Harford, K.; Anjum, A.; Zakir, M.; Wang, F.; Tian, S.; Lee, B.; Liigand, J.; Peters, H.; Wang, R. Q. R.; Nguyen, T.; So, D.; Sharp, M.; da Silva, R.; Gabriel, C.; Scantlebury, J.; Jasinski, M.; Ackerman, D.; Jewison, T.; Sajed, T.; Gautam, V.; Wishart, D. S. DrugBank 6.0: The DrugBank Knowledgebase for 2024. Nucleic Acids Res 2024, 52 (D1), D1265–D1275. 10.1093/NAR/GKAD976.

(14) Lyu, H.; Gharibi, H.; Sokolova, B.; Voiland, A.; Nilsson, B.; Meng, Z.; Gaetani, M.; Saei, A.; Zubarev, R. A. ThermoTargetMiner as a Proteome Integral Solubility Alteration Target Database for Prospective Drugs against Lung Cancer. bioRxiv 2024, 2024.08.06.606599. 10.1101/2024.08.06.606599.

(15) Saei, A. A.; Beusch, C. M.; Chernobrovkin, A.; Sabatier, P.; Zhang, B.; Tokat, Ü. G.; Stergiou, E.; Gaetani, M.; Végvári, Á.; Zubarev, R. A. ProTargetMiner as a Proteome Signature Library of Anticancer Molecules for Functional Discovery. Nat Commun 2019, 10 (1), 1–13. 10.1038/S41467-019-13582-8.

(16) Sabatier, P.; Beusch, C. M.; Saei, A. A.; Aoun, M.; Moruzzi, N.; Coelho, A.; Leijten, N.; Nordenskjöld, M.; Micke, P.; Maltseva, D.; Tonevitsky, A. G.; Millischer, V.; Carlos Villaescusa, J.; Kadekar, S.; Gaetani, M.; Altynbekova, K.; Kel, A.; Berggren, P. O.; Simonson, O.; Grinnemo, K. H.; Holmdahl, R.; Rodin, S.; Zubarev, R. A. An Integrative Proteomics Method Identifies a Regulator of Translation during Stem Cell Maintenance and Differentiation. Nat Commun 2021, 12 (1), 1–16. 10.1038/S41467-021-26879-4.

