## Supplementary Material for "Peptide-to-protein data aggregation using Fisher’s method improves target identification in chemical proteomics"

**Lyu et al.**

**Supplementary Table 1.Drugs and known targets used for FoM calculation**

| **Dataset Name** | **Drug name** | **Targets used for FoM calculation** |
| --- | --- | --- |
| ProtargetMiner_MCF7 deep | Bortezomib | PSMB5, PSMB |
|  | Raltitrexed | TYMS, FPGS |
| OPTI-PISA | MTX | DHFR |
|  | Staurosporine | Kinases:  AAK1, TBK1, TNK2, MAP4K5, MYLK, MAP2K1, CAMK2G, STK24,  CDK2, CAMK2D, PHKG2, CAMKK1, CIT, CHEK1, AURKA,  PDPK1, DAPK3, PIP5K1C, ROCK2, STK10, PAK4, CHEK2,  PRKAR1A, SRC, PRKCA, PRKACA, CSNK2A2, PRKACB,  RPS6KB1, GRK2, MARK3, MAP2K2, CSK, PRKCI, MAPK9,  MAP2K4, MAP2K3, GSK3A, GSK3B, CDK9, RPS6KA3, PRKX,  MAP2K6, MAPK12, PRKAA2, CDK5, CDK16, CDK17, PRKCE,  PTK2, PRKCZ, PRKCD, STK4, PRKAA1, STK3, ROCK1, PRPF4B,  DYRK1A, CAMK1, STK38, RPS6KA1, TAOK1, TLK2, STK32C,  CAMK1D, BRSK2, STK35, PASK, CAMKK2, MKNK1, CDK19,  STK33, SLK, TAOK3, BMP2K, IRAK4, CDK12, NLK, DAPK2,  PKN2, PRKAG1 |
|  | Ganetespib | HSP90AA1, HSP90AA2P, HSP90AB4P, HSP90AA4P, HSP90AB1,  HSP90B1 |
| ThermoTargetMiner | Everolimus | MTOR |
|  | Vorinostat | HDAC1, HDAC2, HDAC3, HDAC6, HDAC8 |
|  | Olaparib | PARP1, PARP2, AKR1C3 |
| OxidoResist_5-FU_PISA-Expression | 5-FU | TYMS |
| HOLSER | MTX | DHFR |
|  | Rapamycin | MTOR, FKBPs (FKBP1A,2,3,4,5,7,8,9,10,11, 14, 15, FKBPL) |
|  | Staurosporine | Kinases:  STK25, GAK, PDPK1, DAPK3, SPAG9, JAK2, RPS6KA4, AK1,  CDK1, PHKG2, FER, PRKCA, PRKACA, PRKACB, RPS6KB1,  JAK1, CDK2, GRK2, MARK3, TYK2, AKT2, TTK, CSK, PRKCI,  GSK3A, RPS6KA3, MAP2K6, PLK1, DAPK1, PRKAA2, CDK5,  PTK2, PRKCD, CDK18, STK4, PAK1, PAK2, STK3, CAMK2G,  MELK, STK38, RPS6KA2, PKN1, PKN2, PKN3, MARK2, MAP4K3,  CAMKK1, PIP4K2C, SLK, IRAK4, MARK1, STK26, TBK1, BAZ1B,  PACSIN3, STK38L, MAP4K5, STK24, ROCK2, WEE1, AXL,  PRKAA1, PRPF4B, CDK9, STK17A, PAK4, CSNK1D, TNK1 |
| AFDIP | MTX | DHFR |
|  | Rapamycin | MTOR, FKBPs (FKBP1A,2,3,4,5,7,8,9,10,11, 14, 15, FKBPL) |
|  | Staurosporine | Kinases:  CAMK2D, CHEK1, PHKG2, YES1, DAPK3, MAP3K7, PRKAR1A,  CDK4, PRKACA, RPS6KB1, CDK9, RPS6KA3, SRPK2, PRKCD,  PRKAA1, STK3, STK38, MARK2, CAMK1D, PIP4K2C, PHKB,  EPS8L2, BMP2K, RIPK2, ROCK2, PAPSS2, CHEK2, PRKAR2A,  ADRBK1, AAK1, PDPK1, MAPK1, MAP2K2, PRKDC, HIPK1, RFK,  CHKB, SPAG9, EPS8, MARK3, TLK2, HGS, SMG1 |

**Supplementary Table 2. Scores of different analysis strategies**

| **Selection of N** | **Protein data** | **Peptide data + Fisher’s method** | | | | | | |
| --- | --- | --- | --- | --- | --- | --- | --- | --- |
|  |  | **All peptides** | **Top N by FC** | | **Top N by p-value** | | **Top N by score** | |
|  |  |  | **no imputation** | **with imputation** | **no imputation** | **with imputation** | **no imputation** | **with imputation** |
| N = 2 | 69.3 | 136.9 | 90.8 | 89.2 | 162.7 | 157.8 | 144.7 | 139.7 |
| N = 3 | 68.5 | 123 | 98.7 | 83 | 161 | 148.1 | 150.9 | 136.7 |
| N = 4 | 67.2 | 126 | 119.6 | 85 | 163.1 | 138.4 | 154 | 132.5 |
| N = 5 | 66.7 | 126.3 | 132.7 | 93.3 | 155.2 | 127.5 | 147.6 | 110.7 |
| N = 6 | 67.6 | 131.9 | 137.6 | 91.4 | 154.5 | 114.6 | 149.2 | 111 |
| *sum* | *339.3* | *644.1* | *579.4* | *441.9* | *796.5* | *686.4* | *746.4* | *630.6* |
